# Identifying the proximate mechanisms that generate variation in nutritional plasticity for fecundity in *Drosophila melanogaster*

**DOI:** 10.1101/2023.03.07.531575

**Authors:** André N. Alves, Avishikta Chakraborty, Mia Wansbrough, Greg M. Walter, Matthew D. W. Piper, Carla M. Sgrò, Christen K. Mirth

## Abstract

Nutrition is an important determinant of an animal’s survival and fitness. Phenotypic plasticity allows a genotype to adjust life history traits to changes in its nutritional environment, and it varies among individuals. To understand how variation in plasticity is achieved, we made use of a *Drosophila melanogaster* isogenic panel to characterize nutritional plasticity for fecundity by feeding flies diets differing in their yeast content and counting the number of eggs produced. We then identified lines with the highest and lowest plastic responses to diet and dissected the potential proximate mechanisms responsible for these differences in plasticity, including morphology, behaviour, and physiology. Our results suggest that genetic variation in plasticity is not due to differences in ovariole number, but due to both increased food intake, and higher efficiency at converting food into eggs. Our results show that, in this population of *D. melanogaster*, variation in behaviour and physiology, but not morphology, underlies differences in plasticity for fecundity. Further, they set the stage for future studies aiming to understand how the proximate mechanisms that generate genetic variation in plasticity contribute to a population’s persistence when faced with environmental changes.

## Introduction

Diet impacts animal wellbeing by shaping a wide variety of life-history traits including growth and development, response to disease, metabolism, lifespan, and reproduction (Simpson & Raubenheimer, 2012; Bozinovic et al., 2011; Pavan et al., 2013; Sgrò et al., 2016). However, not all individuals respond to diet in the same way. The ability of a genotype to adjust its phenotypes in response to environmental conditions, known as phenotypic plasticity, underpins whether a population or species will persist in the face of nutritional stress (Lynch & Walsh, 1998; Hoffmann & Merilä, 1999; Gause, 1947; Bradshaw, 1965; Ørsted et al., 2017). This is particularly important because both the availability and quality of food resources accessible to animals is predicted to vary with climate change (Long et al., 2014; Rosenblatt & Schmitz, 2016). For this reason, understanding how individuals differ in their response to diet, and the underlying causes of these differences, is key to understanding how animals will cope with changes in their food resources.

Genetic variation in plasticity is quantified as genotype-by-environment (G×E) interactions (Schlichting & Pigliucci, 1998; Lande, 2009). Populations residing along environmental gradients respond differently to nutritional stress, suggesting that genetic variation in nutritional plasticity can evolve (Chakraborty et al., 2020, Klepsatel et al., 2020). In addition, individuals within a population show genotype-specific responses to diet (Ng’oma et al., 2018; King et al., 2011, Bergland et al., 2008). Characterising this genetic variation in plasticity is the first step in understanding how populations might respond and eventually adapt to nutritional stress (Narum et al., 2013; Chevin & Hoffmann, 2016; Chakraborty et al., 2020).

While many studies of genetic variation in plasticity within populations have focused on developmental, morphological, behavioural, or stress resistance traits (Van Buskirk & Steiner, 2009; Ghalambor et al., 2007, LaFuente et al., 2018; Cunningham et al., 2020), few studies have directly examined genetic variation in fitness across environments. This is largely because fitness is difficult to measure, and so studies often focus on life-history traits that represent close proxies of fitness, such as lifespan and fecundity (Fisher, 1930). Fecundity is tightly correlated with fitness and is sensitive to changes in nutritional availability and quality (Wallin et al., 1992, Behmer et al., 2001, Lee et al., 2008; Piper et al., 2014), which means small changes in the environment have potential to critically impact fecundity, and therefore fitness. Genetic variation in nutritional plasticity within populations has been documented for fecundity in the fruit fly *Drosophila melanogaster*, measured as the number of eggs laid (Ng’oma et al., 2018; King et al., 2011; Camus et al., 2017). However, little is known about the proximate mechanisms that regulate nutritional plasticity for female fecundity.

Variation in the plastic response of egg laying to nutrition can be regulated by numerous underlying mechanisms, including morphology, behaviour, and physiology. In insects, the ovaries are composed of one or more ovarioles, which are functional units for egg production (David, 1970). For insects like *Drosophila*, the number of ovarioles in an ovary both varies with nutrition and dictates the number of eggs a female can produce at any time (David, 1970). Furthermore, genotypes differ in their plasticity for ovariole number in response to protein restriction (Bergland et al., 2008). Thus, genetic variation in the plastic responses of fecundity across diets could be due to differences in ovary morphology, as determined by ovariole number.

Differences in behaviour, such as rates of food intake, across genotypes can also buffer the extent to which individuals will experience nutritional stress on poor diets. Food intake is carefully regulated to ensure animals reach their nutritional requirements to maximise life-history traits (Simpson & Raubenheimer, 2009; Simpson & Raubenheimer, 1995). Plasticity in food intake and other feeding behaviours occurs when organisms are faced with choices between food sources with different nutritional content or change their intake strategy when forced to eat sub-optimal diets (Carvalho & Mirth, 2017; Rodrigues et al., 2015; Silva-Soares et al., 2017; Diegelmann et al., 2017; Fuhita & Tanimura, 2011). As a result, variation in animals’ feeding behaviours alters whole body physiology, and has the potential to mediate plastic responses in fecundity.

Finally, genetic differences in physiological processes, such as the efficiency with which animals absorb and assimilate the nutrients ingested, can also dictate the degree of plastic response. Traits such as body size have been shown to be regulated by nutrient assimilation (Sibly, 1981; Clissold et al., 2010). Even when animals ingest large quantities of nutrients, they can still develop into smaller adults if they are not able to absorb the nutrients efficiently or are unable to allocate them to the correct organs (Urabe & Watanabe, 1991; Neat et al., 1995). For example, when fed a fixed quantity of the same diet, populations of *D. melanogaster* that are adapted to cold environments reach larger body sizes than those adapted to warm environments (James & Partridge, 1995). Potentially, genetic variation in the ability to absorb and assimilate nutrients could underlie differences in nutritional plasticity for fecundity as well.

One way to study genetic variance in plasticity within a population is to use panels of highly inbred lines originated from natural populations that are exposed to a gradient of environmental conditions. This approach has been used with panels of isogenic lines, such as the *Drosophila melanogaster* Genetic Reference Panel (DGRP) and the *Drosophila* Synthetic Population Resource (DSRP) (Mackay et al., 2012; King et al., 2012). The advantage of using such isogenic lines is that any differences between lines in responses to environmental variation can be attributed to genetic differences in plasticity. Using this approach, studies have revealed genetic variation in plasticity in response to temperature or nutrition for a range of traits such as body size, olfactory behaviour, and wing-body scaling relationships (Lafuente et al., 2018; Sambandan et al., 2008; Frankino et al., 2019).

In this study, we used a newly-derived set of isogenic lines of *D. melanogaster* to quantify within-population genetic variance for nutritional plasticity. We then identified genotypes that showed either high or low plasticity in female fecundity in response to dietary protein. To uncover the source of variation in plasticity, we next explored whether morphology, feeding behaviour, or physiology could explain the differences between high and low plasticity lines. To do this, we counted the number of ovarioles, measured food intake, and assessed protein-to-egg conversion efficiency in these lines across diets varying in protein content. Our study provides insight into the proximate mechanisms that generate within-population genetic variation in plasticity, serving as a platform for understanding how populations can respond to changes in their nutritional environment.

## Methods

### Field collections and establishing experimental lines

We collected 200 field-inseminated *Drosophila melanogaster* females from a banana plantation and Tropical Fruit World in Duranbah, on the east coast of Australia in January 2018 (28.3° S, 153.5° E). We generated two independent isofemale lines from each of the wild-caught females resulting in 400 lines in total. All lines were treated with tetracycline, to attempt to remove *Wolbachia*, and reared for two generations prior to 20 generations of inbreeding (described below). Flies were maintained on standard yeast-potato-dextrose medium (potato flakes 20 g/L; dextrose 30 g/L; Brewer’s yeast 40 g/L; agar 7 g/L, Nipagin 6 mL/L; and propionic acid 2.5 mL/L) at 25 ^o^C, on a 12 h light/dark cycle.

### Generating the isogenic lines

Inbreeding was applied to each of the 400 isofemale lines through full pair-sib mating, for a minimum of 20 generations, resulting in a final panel of 81 fertile isogenic lines which were used for this study (Chakraborty et al., 2023), with an inbreeding coefficient of F=0.986 (Mackay et al., 2012; Reddiex et al., 2018). All lines were maintained at a constant temperature of 18 °C, under a 12 h light/dark cycle, and fed yeast-potato-dextrose medium.

### Fecundity assays

To assess fecundity, flies were maintained for two generations in a common environment of standard yeast-dextrose-potato medium at 25 °C to reduce effects of maternal or grand-maternal environment. To control for larval rearing density and to synchronise adult emergence time, eggs from each isogenic line were collected by allowing adults to lay in embryo collection cages (Genesee Scientific) on 60 mm petri dishes half-filled with apple juice/agar medium, as described in Linford et al., (2013), for 24 h at 25 °C. We then distributed 50 eggs from each line into each of 20 vials containing standard yeast-dextrose-potato food at 25 °C. Adult flies from both sexes that emerged from these vials were collected over a 48 h period, transferred to new vials containing yeast-dextrose-potato medium, and left to interact for 48 h, during which they could mate multiple times, prior to the assessments of fecundity described below.

After 48 h of mating, females were separated from males, and five females from each isogenic line were transferred into vials that contained one of three diets, to assess nutritional plasticity in fecundity. To test for changes in fecundity due to protein restriction, we compared a 100% standard diet (yeast-dextrose-potato) with two other diets that contained 5% or 50% of yeast (relative to the 100% diet), but the same amount of dextrose and potato. We chose these diets because yeast is the main source of protein for *D. melanogaster*, and protein concentration correlates tightly with egg production (Piper et al., 2014; Ortega et al., 2021; Alves et al., 2022). The low protein diets chosen for this study simulates not only variation in diet that animals already experience (Markow & O’Grady, 2008; Matavelli et al., 2015; Silva-Soares et al., 2016), but also the predicted reductions in nutrition expected under climate change (DaMatta et al., 2010; Sardans et al., 2017).

To quantify changes in fecundity for the three diets, we established ten replicate vials, each containing five females from each isogenic line for each diet/line combination. Females were transferred to fresh food every 24 h. Eggs laid on days 5, 6 and 7 of the assays (days post-mating) were counted since this interval is known to capture the peak period in fecundity for *D. melanogaster* and is often used as a measurement for fecundity (Novoseltsev et al., 2003).

### Testing for overall genotype-by-environment interactions for fecundity

To test for significant genotype-by-environment interactions for fecundity, we applied a generalised linear mixed-effects model using a Poisson distribution with fecundity (the sum of eggs laid from day 5 to 7) as the response variable. To apply the linear model, we used the glmmTMB package in R. We fit sequential models (Table 1) with diet as a fixed effect and experimental block as a random effect. Experimental block was defined as a set of isogenic lines tested over the same period of time, as it was physically impossible to test all 81 lines concurrently. To test for genotype-by-environment interactions, we used random regression. This tested for improved fit between models in which only block was including as random effect, block and isogenic line were included as a random effects, and block and the interaction between diet and isogenic lines were included as random effects (Table 1). We tested for improved fit using a likelihood ratio test and by comparing AIC values across models (Table 1).

**Table 1:**
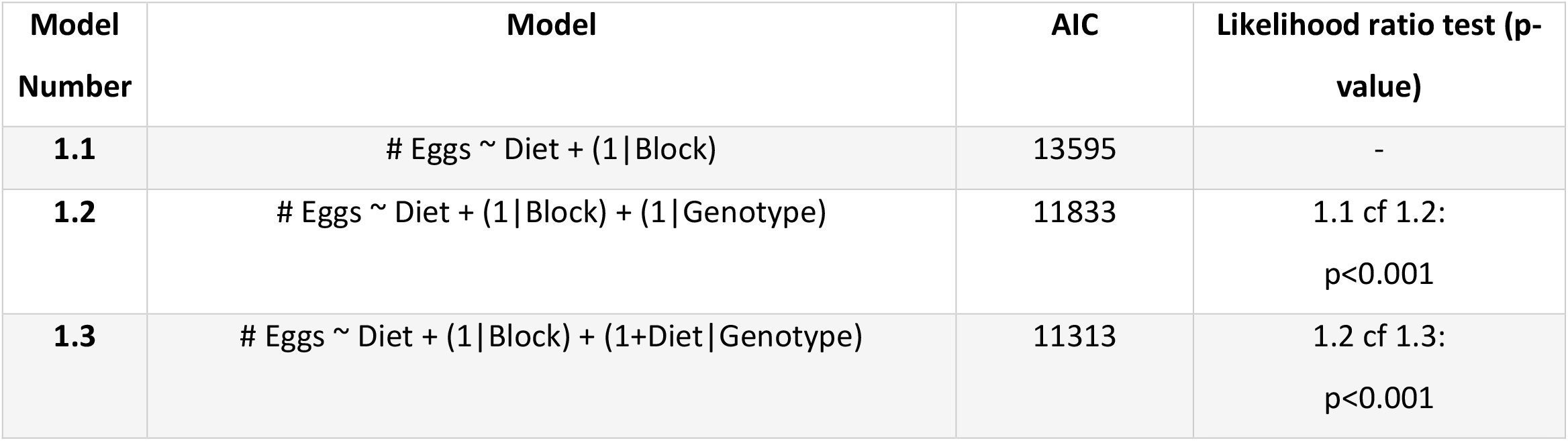
Random regression mixed effects models testing for genetic variation in plasticity for fecundity in response to the yeast content of the diet. Data was fit using a Gaussian distribution.

### Quantifying genetic variance in fecundity

We then explored whether G×E across diets was the result of changes in genetic variation for fecundity across diets or due to genetic correlations across diets of less than 1 (Falconer 1952; Via & Lande, 1985). We used Bayesian models to estimate genetic variance within environments, and genetic correlations in fecundity between diets using a generalized linear mixed model with the R package ‘MCMCglmm’ (Hadfield, 2010):

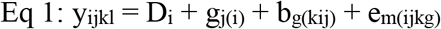

where the sum of eggs laid from day 5 to 7 was included as a univariate Gaussian-distributed response variable (y_ijkl_). The *i*th diet was included as the only fixed effect (D_i_). The *j*th isogenic line (g_j(i)_) and experimental block (b_l(kij)_) were included as random effects to account for variation among lines within each diet, and among experimental blocks. Residual variance is represented by e_m(ijkg)_. The model was implemented with 1.1 million iterations, a burn-in of 100,000 iterations, and a thinning interval of 500 iterations. We checked autocorrelation for model convergence by confirming that the effective sample size exceeded 85% of the number of samples that were saved. For each diet we estimated random intercepts and slopes, which calculated among-genotype genetic variance within each diet as well as the genetic covariance between diets, resulting in a 3×3 genetic covariance matrix.

To quantify the genetic correlation between fecundity across environments, a correlation matrix was calculated from the covariance matrix of fecundity. To consider the uncertainty in our estimates of genetic variance and genetic correlations between treatments, we calculated the 95% Highest Posterior Density (HPD) credible intervals. Density distributions of genetic variance and genetic correlations were created by extracting 2,000 samples from the model.

### Identifying lines with high/low plasticity in fecundity

To characterise differences in plasticity across genotypes, we first eliminated any isogenic lines that had laid less than three eggs over the time period measured across all diets as we could not accurately characterise plasticity in these lines (Figure 2A). To obtain coefficients of plasticity across the remaining isogenic lines, we calculated the normalised difference in egg numbers by subtracting the number eggs laid on the 5% from those laid on the 100% yeast content diets for each replicate/genotype and normalising this difference by the mean egg number in the 100% diet for each genotype (Figure 2B). We then ran linear mixed effect models on the normalised difference in egg numbers using the genotype as a fixed effect and block as a random effect (‘lme4’ package). We calculated the estimated marginal means for each genotype using the ‘emmeans’ package. Finally, we selected five of the ten genotypes with the highest emmeans values as the high plasticity lines, and five out of the ten lines with the smallest emmeans values, as the low plasticity lines (Figure 2C). The lines characterised for their high or low plasticity chosen from the analysis were maintained for two generations in common garden settings of standard yeast-dextrose-potato medium at 25 °C prior to all assays described below.

**Figure 1:**
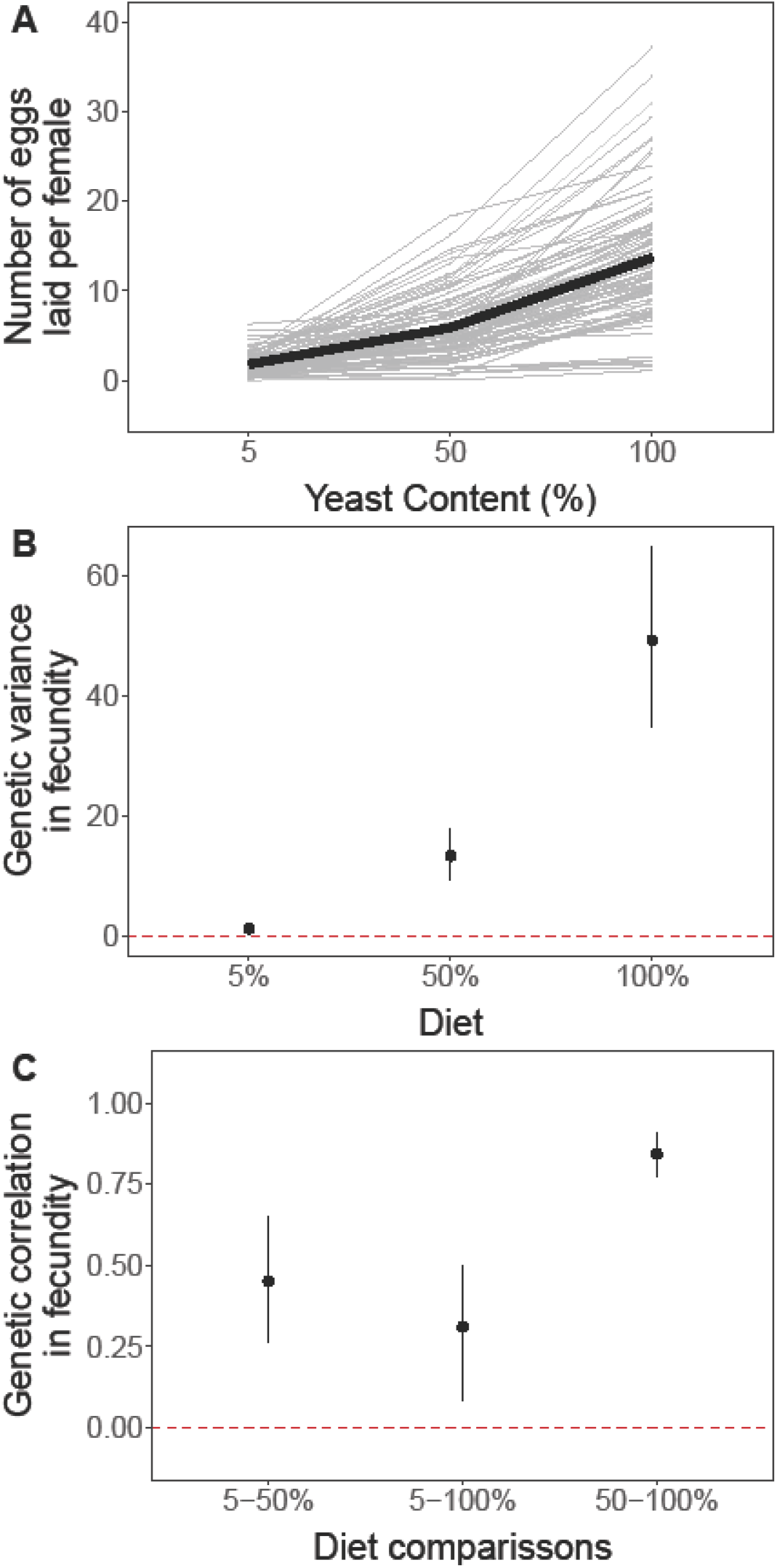
Nutritional plasticity for fecundity and its underlying variation. A) Reaction norms of fecundity for each isogenic line across diets with 5, 50 and 100% yeast content. The sum of eggs laid between days 5 to 7 for each female were counted in each diet for each isogenic line. The mean reaction norm of the population is represented by the black line and each isogenic line is represented in grey. B) Estimate of genetic variance in fitness and respective 95% credible interval for each diet. The dashed line represents 0 genetic variance, i.e. no variation within that diet. C) Estimates of genetic correlation and respective 95% credible intervals. The dashed line represents 0 genetic correlation, i.e. no genetic correlation between diets.

**Figure 2:**
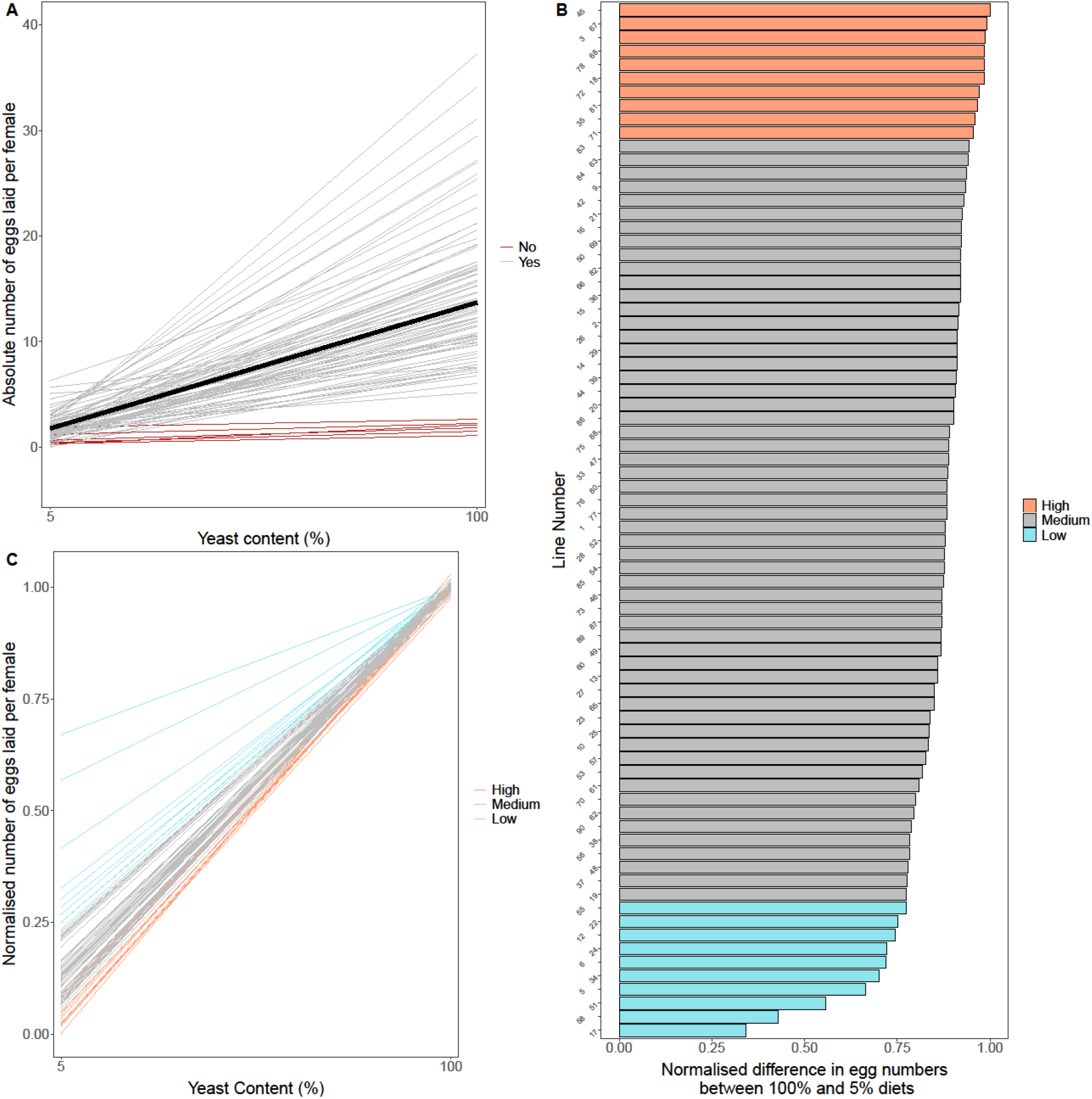
Characterizing plasticity across isogenic lines. A) Isogenic lines excluded from further analysis. Lines in red represent the isogenic lines that laid less than 3 eggs across all diets and were excluded from further analysis. B) Ranking isogenic lines by Best Linear Unbiased Predictors (BLUP) value on the normalized difference between the number of eggs laid on 100% and 5% diets. Blue lines show low plasticity lines with low normalized difference between egg numbers between 100% and 5% diets. Orange lines represent high plasticity lines with a high normalized difference between egg numbers between 100% and 5% diets. C) Representation of reaction norms for each isogenic line. Normalised number of eggs per female was plotted for both diets selected, with blue lines representing the low plasticity lines, which have shallower slopes, and orange lines representing high plasticity lines, which have steeper slopes.

### Ovariole dissection and counting

To assess if the differences in plasticity in fecundity between the high and low plasticity lines arose from differences in the number of ovarioles, flies were reared at 25°C in standard potato-yeast-dextrose medium at controlled densities of 50 eggs per vial of food with 3 replicate vials per line. Once eclosed, adult flies were collected over a 48-hour period, transferred to new vials containing standard yeast-dextrose-potato medium, and left to mate for 48 h. At the end of this period, female flies were transferred from three vials per line into two microtubes, placed in dry ice and stored at −80°C until ready for dissection.

Ovariole number was measured on 10 females per replicate microtube (20 females per line). Flies were submerged in 1 × phosphate-buffer solution (PBS) and their ovaries removed and teased apart to count the number of ovarioles.

### Food intake and fecundity assays

We next examined the relationship between food intake and fecundity in the high and low plasticity lines. To control for larval rearing density and to synchronise adult emergence time, we collected eggs from the parental generation by leaving them to lay in embryo collection cages (Genesee Scientific) on 60 mm petri dishes half-filled with apple juice/agar medium, as described in Linford et al. (2013), for 24 h at 25°C. Eggs were then transferred into 10 food vials per line, at a density of 50 eggs per vial, each of which contained 6 mL of standard yeast-dextrose-potato medium at 25°C.

The adult flies that emerged from these vials were collected over a 48-hour period, transferred to new vials containing standard yeast-dextrose-potato medium, and left in vials for 48 h to allow the flies to mate multiple times. Following the 48 h mating period, five female flies per line were transferred into vials that contained 3 mL of apple juice/agar medium to a total of 10 vials per line and per diet, as described below.

We used the experimental setup for the capillary-assisted feeding (CAFE) assays from Diegelmann et al. (2017) to measure food intake and number of eggs laid. We pierced narrow cotton flugs (Genesee Scientific, product code 076-49-103) three times with a sharp, hot metal rod of the same diameter as the calibrated glass micropipettes used for the assays (5 µL, Sigma, product number P0549). Three glass micropipettes per vial were filled with liquid media, marked with a black marker at the meniscus, and inserted in the cotton flugs. The media used consisted of either 5N or 100N holidic food without agar (a synthetic diet containing the proportion of each amino acid according to an exome-matching study (Piper et al., 2014)). In these diets N represents the amount of nitrogen available, and by association, amount of protein available in the food, with 100N equal to 100 mM of nitrogen available and 5N, 5 mM of available nitrogen (Piper et al., 2014). This medium was used over a standard yeast-dextrose-potato medium since the former is soluble, whereas the latter settles to the bottom of the capillary, impeding flies from feeding from it. We also added 0.1% blue food colouring to the media for contrast when measuring food intake (Queen food dye, blue, batch #118106). We established ten replicate vials per line and diet combination, each containing five females.

Concurrently, to measure number of eggs laid each day, females from the capillary feeding vials were transferred to fresh apple juice/agar vials every 24 h for seven days, after which the eggs laid over the previous 24 h period were counted manually. Capillaries were replaced, and amount of food left in the capillary from the previous 24 h was measured with the same frequency to obtain feeding intake measurements. This was done for seven consecutive days. The CAFE setup was placed inside a plastic storage box (25 cm (H) x 48 cm (W) x 34.5 cm (D)) containing wet paper towel and closed with a lid to avoid evaporation. The CAFE setup was placed in an incubator under a 12 h light/dark cycle kept at 25 °C and 80% humidity. Apple juice/agar vials containing media-filled capillaries, but no flies, were placed in the same boxes to measure evaporation of the media. Evaporation losses were subtracted from experimental readings to obtain true ingestion amounts (Diegelmann et al., 2017).

### Assessing statistical significance across proximate mechanisms

To assess differences in ovariole number across plasticity lines, we fit the data with a linear mixed effects model using the ‘lme4’ package. The average number of ovarioles/ovary was used as a response variable, and plasticity group was used as a fixed effect. Microtube and isogenic line were included as random effects.

Next, we tested for significant differences in egg laying behaviour between high and low plasticity lines. A generalised mixed effects linear model was created with the sum of eggs laid throughout the experiment as a Poisson-distributed variable with a log link function. Diet and plasticity group were included as fixed effects. Experimental block, replicate, storage box, and isogenic line were included as random effects.

To test if the amount of ingested food differed between high and low plasticity lines, food intake over the length of this experiment was calculated for each food/line pairing. Since the food intake data was not normally distributed, we square-root transformed the data, which improved the fit. The transformed food intake was used as a response variable and diet and plasticity group were included as fixed effects. Block, replicate, evaporation box, and isogenic line were included as random effects.

To test if protein ingested was a significant predictor of female fecundity, we used a generalised mixed effects linear model as described above. The sum of eggs laid throughout the experiment was the response variable, which was transformed using a log(10) scale. The same transformation was applied for protein ingested, which was calculated considering the amount of nitrogen available in the food. Plasticity group, and protein ingested were included as fixed effects. Block, storage box, and isogenic line were included as random effects.

All statistical analyses were performed in R Studio (version 3.4.1). Plots were produced using ggplot2 (‘tidyverse’ package).

## Results

### Testing for within-population genotype-by-environment interactions for fecundity

To test for overall genetic variance in the plastic response of fecundity to nutrition, we placed isogenic lines onto diets differing in yeast content and counted the number of eggs laid on days 5-7. We found that total fecundity significantly increased with yeast content in the diet, and that genotypes differed in the total number of eggs laid across all diets (Figure 1A). Furthermore, genotypes differed in their response to diet, resulting in a significant diet by genotype interaction. Including a genotype by diet interaction in the random effects significantly improved the model fit over models that included either block alone or block and genotype, as indicated by a significant likelihood ratio test and a lower AIC score (Table 1). Thus, we observed significant a genotype-by-environment (GxE) interaction for egg numbers.

### Quantifying genetic variance in the plastic response of fecundity to diet

Having detected significant GxE for fecundity, we next determined the origin of this variation. G×E can be due to increased variation in fecundity on specific diets across genotypes (Sheth et al., 2018). Alternatively, G×E can result from differences in the response of genotypes to diets, resulting in genetic correlation of less than 1 across diets.

We found significant genetic variance for fecundity on all three diets, and the expression of genetic variance increased with yeast concentration, as indicated by non-overlapping credible intervals and density distributions (Figure 1B, Supplementary Figure 1). Thus, changes in genetic variance across diets explains at least some of the significant G×E interaction for fecundity.

Genetic correlations in fecundity among diets were all strong and positive (Figure 1C, Supplementary Figure 1). Genetic correlations were highest between the 50% and 100% yeast diets, and significantly lower between the 5% and 50% and 5% and 100% diets. The fact that genetic correlations were less than one for all three diet comparisons indicates that the significant genotype-by-environment interaction for fecundity also results from the fact that the isogenic lines differ in some of the alleles that underpin nutritional plasticity for this trait.

### Characterising plasticity across isogenic lines

We next characterised differences in plasticity for female fecundity across low (5%) and high (100%) dietary protein concentrations. We first excluded any isogenic lines that laid less than 3 eggs across the time period observed for all diets as we could not accurately characterise plasticity in these lines (Figure 2A). With the remaining lines, we calculated the estimated marginal means (emmeans), using the normalised difference between the 5% and 100% yeast content diets for each genotype (Figure 2B). These means estimate how genotypes rank relative to each other and the mean. We then selected five of the ten genotypes with the smallest emmean values as the high plasticity lines, and five out of the ten lines with the largest emmean values, as the low plasticity lines (Figure 2B,C). We then used these chosen lines to explore three proximate sources of variation in plasticity for fecundity: 1) ovary morphology, measured by ovariole number, 2) feeding behaviour, and 3) physiology, determined by the efficiency at which genotypes convert ingested protein into eggs.

### Measuring ovariole number across plasticity lines

Ovariole number is known to limit the number of eggs produced (David, 1970), but it is unclear whether it plays a role in plastic responses of egg laying to diet. We hypothesised that isogenic lines that have capacity to lay more eggs in the most protein rich diets might have higher ovariole number, and vice-versa. To test whether plastic responses were determined by ovariole number, we reared our high and low plasticity isogenic lines on the same media and dissected groups of females, teasing apart their ovaries to count ovarioles. The number of ovarioles did not differ between plasticity groups (Figure 3A, Supplementary Table 1). These data indicate that differences seen in nutritional plasticity for fecundity are not due to differences in ovariole number between isogenic lines.

**Figure 3:**
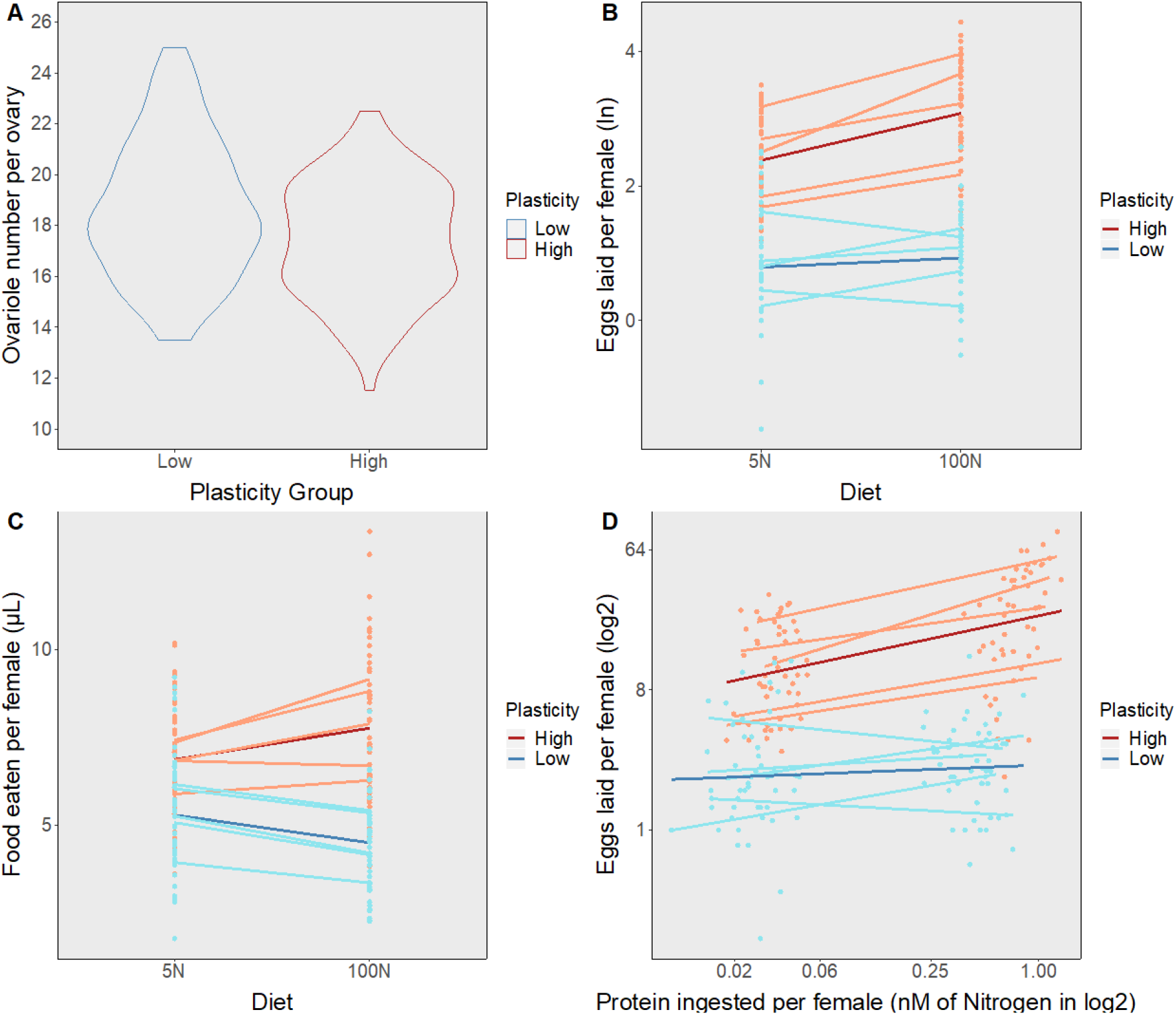
The relationship between differences in plasticity across isogenic lines, ovary size, food intake, and the capacity to convert food into eggs. A) Ovariole number in isogenic lines with different levels of plasticity. Number of ovarioles per ovary in flies from low (cadet blue) or high (light salmon) plasticity groups. Points represent data from each isogenic line/microtube. B) Eggs laid per female in diets containing 5% (5N) or 100% (100N) protein across isogenic lines with different levels of plasticity. C) Food intake per female in diets containing 5% (5N) or 100% (100N) protein across isogenic lines with either high or low levels of plasticity. D) The relationship between number of eggs laid and amount of protein ingested in high and low plasticity lines subjected to diets differing in protein content and allowed to lay eggs for 7 days. Points represent data from each replicate vial/diet/line. Salmon coloured points and lines are high plasticity lines and cadet blue lines are low plasticity lines. Darker, bold lines with standard error represent the average for each plasticity group.

### Characterising feeding behaviour between high and low plasticity lines

Differences in plasticity might also arise due to differences in food intake between high and low plasticity lines in response to differing diets. Female flies from each of the high and low plasticity lines were placed in vials with access to three glass microcapillaries filled with holidic medium containing either 5% or 100% of the total protein of the standard holidic diet (Piper et al., 2014), allowing us to quantify the total amount of protein consumed.

As expected, the number of eggs laid was influenced by the protein concentration in the capillaries (Figure 3B, Table 3, Supplementary Table 2), but this depended on the plasticity grouping. On average, high plasticity lines laid five to nine more eggs than the low plasticity lines across both diets (Supplementary Table 2). In addition, high plasticity lines showed a steeper response in the number of eggs laid in response to the protein concentration in the diet than the low plasticity lines, resulting in a significant interaction term between diet and plasticity group (Figure 3B, Table 3). Furthermore, the fact that these lines retained either high or low plasticity on the holidic medium (this experiment) in addition to that on yeast-based food (Figure 1A) suggests that this grouping method is accurate to ascertain plasticity across diet types and lines.

**Table 3:**
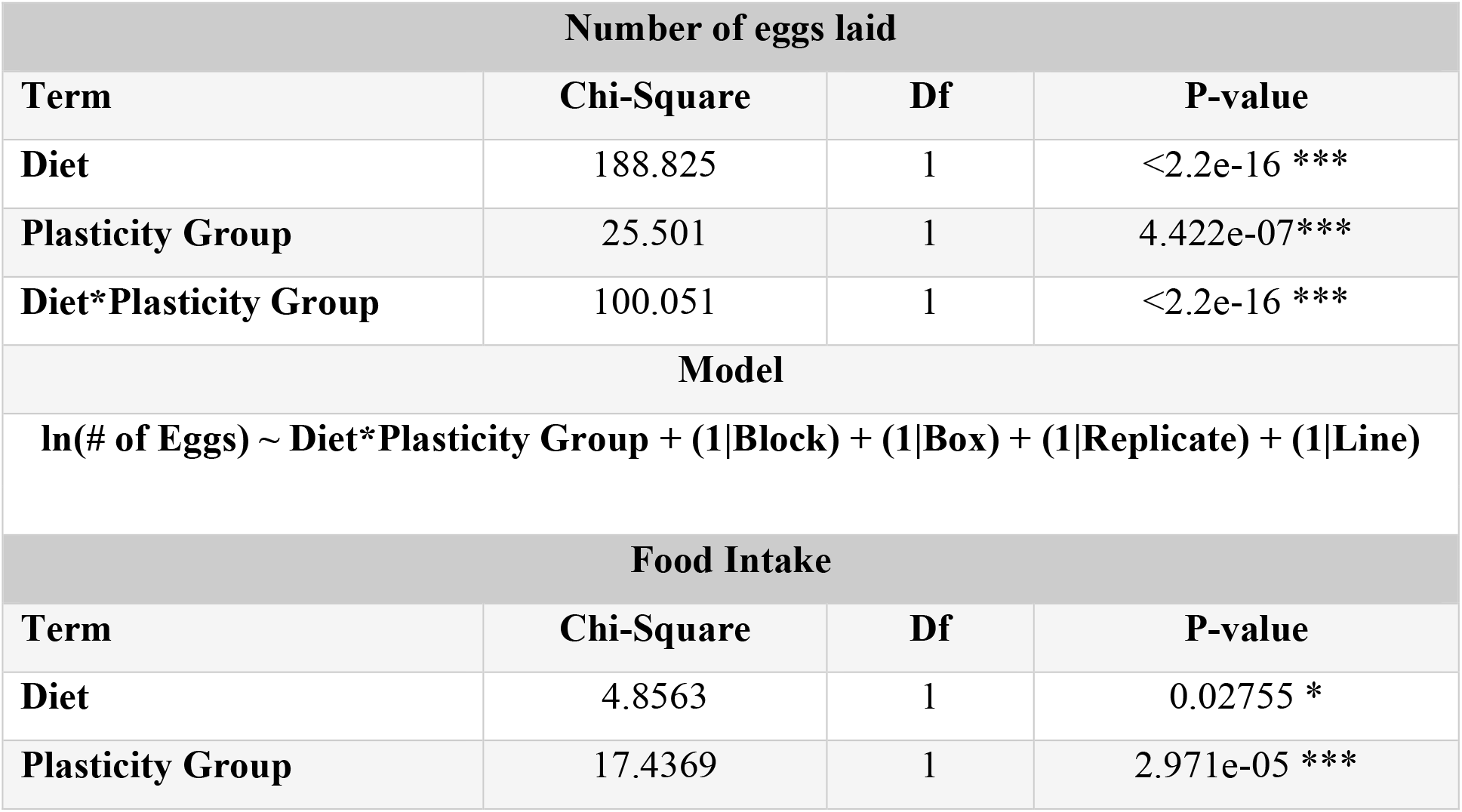
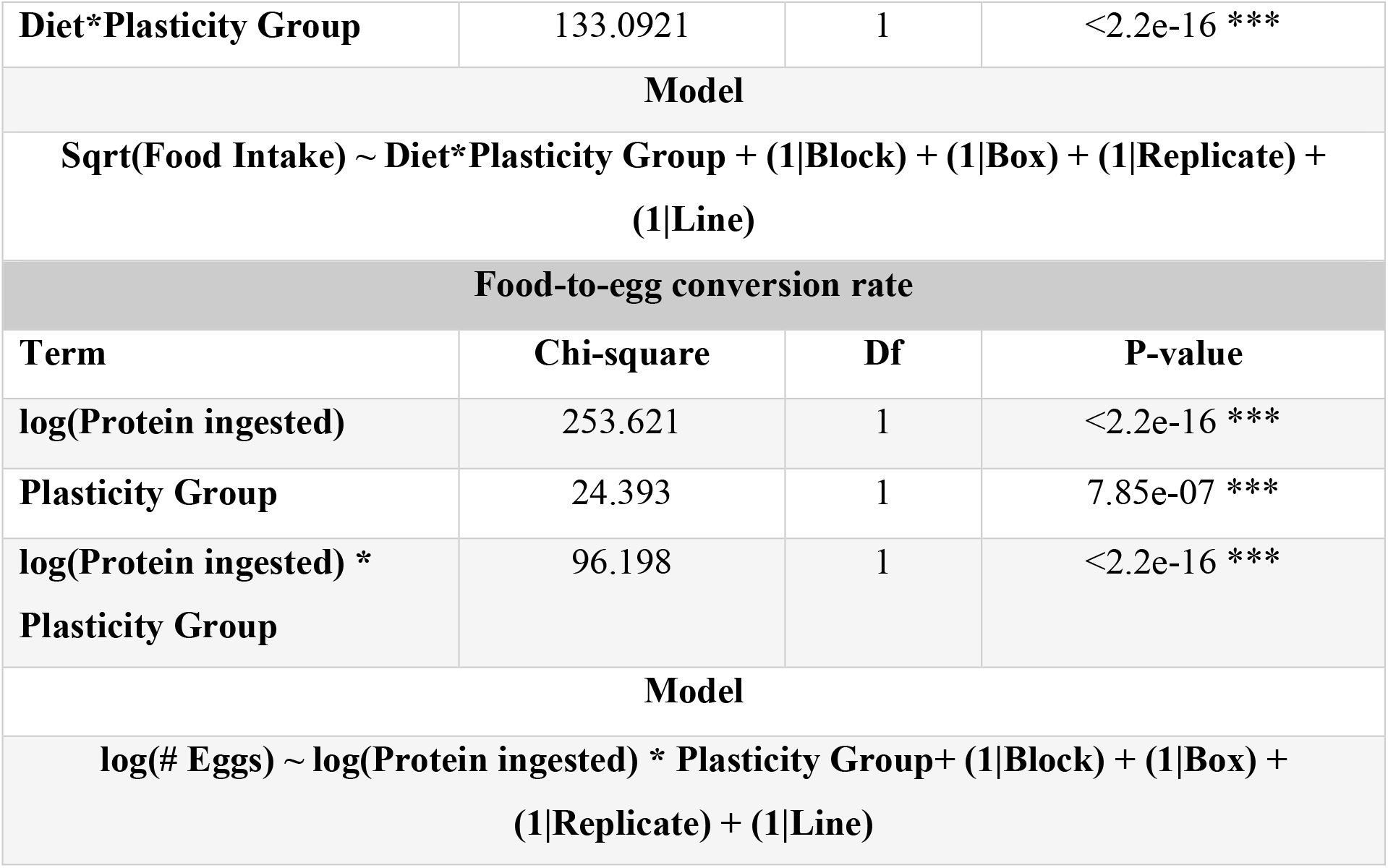
Manipulating both diet and plasticity groups (high or low plasticity) alters egg laying or food ingestion behaviours. Egg laying data was fit with a generalized mixed effects model, whereas ingestion was fit with a linear mixed effects model. A generalized mixed effects model was fit to test the effect of diet, food ingestion and plasticity on egg laying.

| Number of eggs laid |  |  |  |
| --- | --- | --- | --- |
| Term | Chi-Square | Df | P-value |
| Diet | 188.825 | 1 | <2.2e-16 *** |
| Plasticity Group | 25.501 | 1 | 4.422e-07*** |
| Diet*Plasticity Group | 100.051 | 1 | <2.2e-16 *** |
| Model |  |  |  |
| $\ln(\# \text{ of Eggs}) \sim \text{Diet*Plasticity Group} + (1\vert \text{Block}) + (1\vert \text{Box}) + (1\vert \text{Replicate}) + (1\vert \text{Line})$ | | | |
| Food Intake |  |  |  |
| Term | Chi-Square | Df | P-value |
| Diet | 4.8563 | 1 | 0.02755 * |
| Plasticity Group | 17.4369 | 1 | 2.971e-05 *** |
| **Diet\*Plasticity Group** | 133.0921 | 1 | <2.2e-16 *** |
| **Model** |  |  |  |
| **Sqrt(Food Intake) ~ Diet\*Plasticity Group + (1\|Block) + (1\|Box) + (1\|Replicate) + (1\|Line)** |  |  |  |
| **Food-to-egg conversion rate** |  |  |  |
| **Term** | **Chi-square** | **Df** | **P-value** |
| **log(Protein ingested)** | 253.621 | 1 | <2.2e-16 *** |
| **Plasticity Group** | 24.393 | 1 | 7.85e-07 *** |
| **log(Protein ingested) \* Plasticity Group** | 96.198 | 1 | <2.2e-16 *** |
| **Model** |  |  |  |
| **log(# Eggs) ~ log(Protein ingested) \* Plasticity Group + (1\|Block) + (1\|Box) + (1\|Replicate) + (1\|Line)** |  |  |  |

The concentration of protein in the diet also had a significant impact on food intake (Table 3). The high plasticity lines ate significantly more food than the low plasticity lines, and this difference was more apparent on the high protein diet (Figure 3C, Table 3, Supplementary Table 2), averaging 3.2 µL more food on a high protein diet. Furthermore, the high plasticity lines showed a mild increase in food intake with increasing protein content of the diet, while the low plasticity lines reduced their food intake slightly with increasing dietary protein concentration, resulting in a significant interaction between diet type and plasticity group for food consumption (Figure 3C, Table 3). This suggests that high and low plasticity lines differ in the way they regulate their food intake in response to the protein content of the diet.

### Measuring differences in food-to-egg conversion rates between low and high plasticity lines

To understand if variation in egg laying plasticity was due to the amount of food eaten or ability to utilise nutrients more efficiently, we next examined the relationship between amount of protein ingested and the number of eggs laid across plasticity groups. In general, the number of eggs laid increased with protein ingested, suggesting that amount of protein ingested is a reasonable predictor of fecundity (Figure 3D, Table 3). The interaction between protein ingested and plasticity group was also significant, showing that high and low plasticity lines vary in the relationship between the number of eggs laid and the amount of protein ingested (Figure 3D, Table 3). The number of eggs laid increased at a faster rate with each nM of nitrogen ingested in high plasticity lines when compared to low plasticity lines for the same amount of protein ingested. This difference in conversion rate is seen as a higher slope for the high plasticity lines (Figure 3D, Table 3, Supplementary Table 2). These results suggest that food intake alone cannot explain the difference in plasticity between high and low plasticity lines, and that other mechanisms such as food absorption and/or assimilation also contribute to the response.

## Discussion

Nutritional stress is one of the most common stressors faced by animals in nature and is likely to become increasingly so with climate change (Scheffers et al., 2016; Houghton et al., 1990; Dawson et al., 2011; Thomas et al., 2004; Foden et al., 2013). Phenotypic plasticity will play a key role in shaping how animals will respond to climate change, at least in the short to mid-term (Lynch & Walsh, 1998; Hoffmann & Merilä, 1999; Franks et al., 2007; Geerts et al., 2015). Genetic variation in plasticity exists both across (Chakraborty et al., 2020, Klepsatel et al., 2020) and within (Schlichting and Pigliucci 1998; Lande 2009; LaFuente et al., 2018; Chakraborty et al., 2022; Dick et al., 2011) populations. Moreover, modulating the plastic response of life-history traits to nutrition is vital for an organism to persist through changes in their environment. Despite its importance, the relative contribution of variation in the proximate mechanisms that regulate plastic responses to diet is poorly understood. Here, we used isogenic lines of *D. melanogaster* to analyse the source of genetic variation for egg laying plasticity, and to understand how ovariole number, food intake, and protein-to-egg conversion efficiency impacted fecundity, a key trait correlated with fitness.

### The source of genotype-by-environment interactions in isogenic lines

Numerous studies have revealed within-population genetic variation for plasticity in a range of life history traits (Van Buskirk & Steiner, 2009; Ghalambor et al., 2007, LaFuente et al., 2018; Cunningham et al., 2020, Dick et al., 2011), but only a few have assessed plasticity for fecundity, an important proxy for fitness (Ng’oma et al., 2018; King et al., 2011; Camus et al., 2017). Such studies emphasise that the distribution of reaction norms for individual genotypes can vary markedly from the population-level reaction norm. Our current study adds to the existing body of work by revealing that individual genotypes within a population can vary not only in the distribution of reaction norms for individual genotypes, but also in the extent and direction of plasticity for fecundity.

We then went on to reveal the cause of this within-population G×E. Significant G×E can arise as the result of differences in the expression of genetic variation across environments and/or cross-environment genetic correlations that are less than one (Falconer 1952; Via & Lande 1985; Sheth et al., 2018). While some studies have found that exposure to environmental stress may increase the expression of genetic variance for life-history traits, including fecundity (Service & Rose, 1985; Van Noordwijk et al., 1988; Etges, 1993; Holloway et al., 1990), other studies suggest that this might depend on the trait and the environmental condition studied (Larsson et al., 1997; Jenkins et al., 1997; Sgrò & Hoffmann, 1998). We found significant levels of genetic variance in fecundity in all three nutritional environments, however the expression of genetic variance increased with increasing diet quality. In this case, stressful diets reduced genetic variance instead of increasing it.

Although several theoretical and experimental studies have shown that genetic correlations can change in sign and magnitude when individuals are exposed to new or stressful environments (Krebs & Loeschcke, 1994; Norry & Loeschcke, 2002; Sgrò & Hoffmann, 2004), we found that all three cross-diet genetic correlations were large and positive, which is consistent with other studies that show positive genetic correlations across environments (Ebert et al., 1993, Etges, 1993, Windig, 1994). The significant G×E we found for fecundity can be explained by both changes in genetic variance across diets and the fact that genetic correlations were less than one, indicating that the alleles that contribute to fecundity are in part independent across environments. Alleles that contribute to fecundity independently across environments might, therefore, explain differences in plasticity across different genotypes.

### Proximate mechanisms that regulate plasticity

Ovariole number is a significant predictor of fecundity in *Drosophila*, as it caps the maximum number of eggs that can be produced in optimal conditions (David, 1970). Differences in ovariole number have been studied extensively between *Drosophila* species, and it is known that environmental factors such as nutrition and temperature regulate this number (Markow & Grady, 2008; Hodin & Riddiford, 2000; Bergland et al., 2008). Genetic differences in ovariole number result from processes occurring during larval development, when the ovarian structures first develop (Sarikaya et al., 2012). We hypothesized that genetic differences in these developmental processes might lead to differences in ovariole number, thereby limiting plasticity in fecundity. However, our data show that variation in plasticity for fecundity in this population is unlikely to arise due to differences in ovariole number.

Previous studies have shown that flies regulate their food intake by choosing to eat different diets to maximise life-history traits. For instance, flies eat salt-rich diets to increase fecundity (Walker et al., 2015), diets with intermediate protein to carbohydrate ratios to minimise development time (Rodrigues et al., 2015), and diets with lower protein to carbohydrate ratios to modulate survival to infection (Ponton et al., 2019). Our study highlights those differences in food intake correlate with differences in nutritional plasticity in fecundity as well.

However, variation in food intake does not account for all the differences in nutritional plasticity in fecundity observed across the high and low plasticity lines in the current study. Especially with the low protein diet, we found that high and low plasticity lines showed little difference in food intake, despite significant differences in the number of eggs laid. This suggests that other post-ingestive, physiological mechanisms, such as amino acid absorption or nitrogen retention, underlie genetic variation in this trait. Previous studies using cabbage white, *Pieris rapae*, larvae that fed on different plants showed that larvae differ in their amino acid absorption depending on the diet they are fed (Slansky & Feeny, 1977). Variation in the efficiency of nutrient absorption and/or assimilation can also impact growth, development, and longevity (Patt et al., 2003; Min et al., 2007). Thus, variation in the efficiency with which animals can convert nutrients into phenotypic outputs is common.

A complementary, powerful approach that could provide further insight into the proximate mechanisms that generate differences in plasticity in fecundity would be using Genotype Wide Association Studies (GWAS) in isogenic panels such as the *Drosophila* Genetic Reference Panel (DGRP) to identify genes that contribute to differences in food intake and nutrient assimilation. This approach has been used successfully to identify loci contributing to variation in thermal plasticity in body size (LaFuente et al., 2018). Other studies have found genes associated with traits such as starvation resistance, olfactory behaviour, and body mass composition that increase variation in plasticity when genotypes are exposed to different diets (Nelson et al., 2016; Sambandan et al., 2008). A SNP affecting mean food intake has already been discovered through this approach (Garlapow et al., 2015). An approach like this could be applied to the isogenic panel used for this study to uncover new genes associated with the differences in intake or nutrient utilization observed.

To further understand plasticity, it would be of great value to examine plasticity in male fitness in these isogenic lines. Differences in this trait could be a by-product of sexually dimorphic gene expression. Previous studies have uncovered genetic variation in male and female dietary requirements, which leads to differences in feeding behaviour between the two sexes (Camus et al., 2017). In isogenic lines, gene expression might be severely skewed towards one sex across diets, which can lead to a detrimental gene expression for the other sex (Connallon & Clark, 2011). This could potentially lead to differences in plasticity, and using GWAS studies, we could uncover loci associated with sexually dimorphic gene expression and associated with variation in plasticity.

Variation in plasticity can be achieved in several ways. Here we show that variation in nutritional plasticity for fecundity is not explained by morphological differences in ovariole number, but due to differences in food intake and the ability to convert ingested nutrients into eggs. Additional research into the potential loci responsible for creating these differences will provide insight into the genetic mechanisms underlying variation in plasticity. Furthermore, this study enhances the importance of understanding the proximate mechanisms that generate variation in plasticity, since these will regulate the fitness-related traits that will ultimately contribute to species’ persistence and adaptation.

## Acknowledgements

We would like to thank members of the Mirth, Sgrò, and Piper labs, past and present, for their help throughout this project. A special thank you to Tahlia Fulton for their unrelenting help in CAFE assays. We would also like to thank Brendan Houston for their help cleaning and refilling capillaries for the CAFE assays. This research was funded by ARC (DP180103725) for CMS, ARC (FT170100259) for CKM, and ARC (FT150100237) and NHMRC (APP1182330) for MDP.

## Author contributions

ANA contributed with experimental design, execution, data analysis and interpretation. AC contributed with experimental design and execution and data analysis. MW contributed with experimental design and execution. GMW contributed with data analysis and interpretation. MDW, CMS and CKM contributed with experimental design, and data analysis and interpretation. ANA, AC, GMW, MDW, CMS, and CKM contributed to writing and revising the manuscript.

## Data availability

All data and scripts are available in Figshare (DOI: 10.26188/28079516).

## Supplementary Figures and Tables

**Supplementary Table 1:**
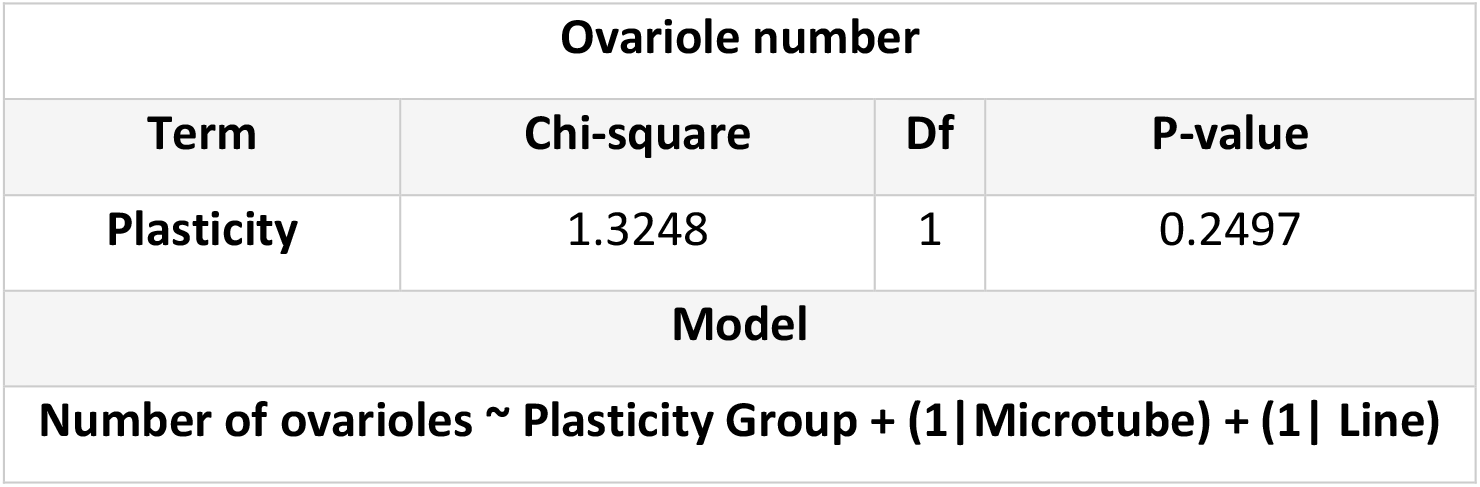
Ovariole number does not differ among plasticity groups (high or low plasticity). Ovariole number was fit with a linear mixed effects model.

**Supplementary Table 2:**
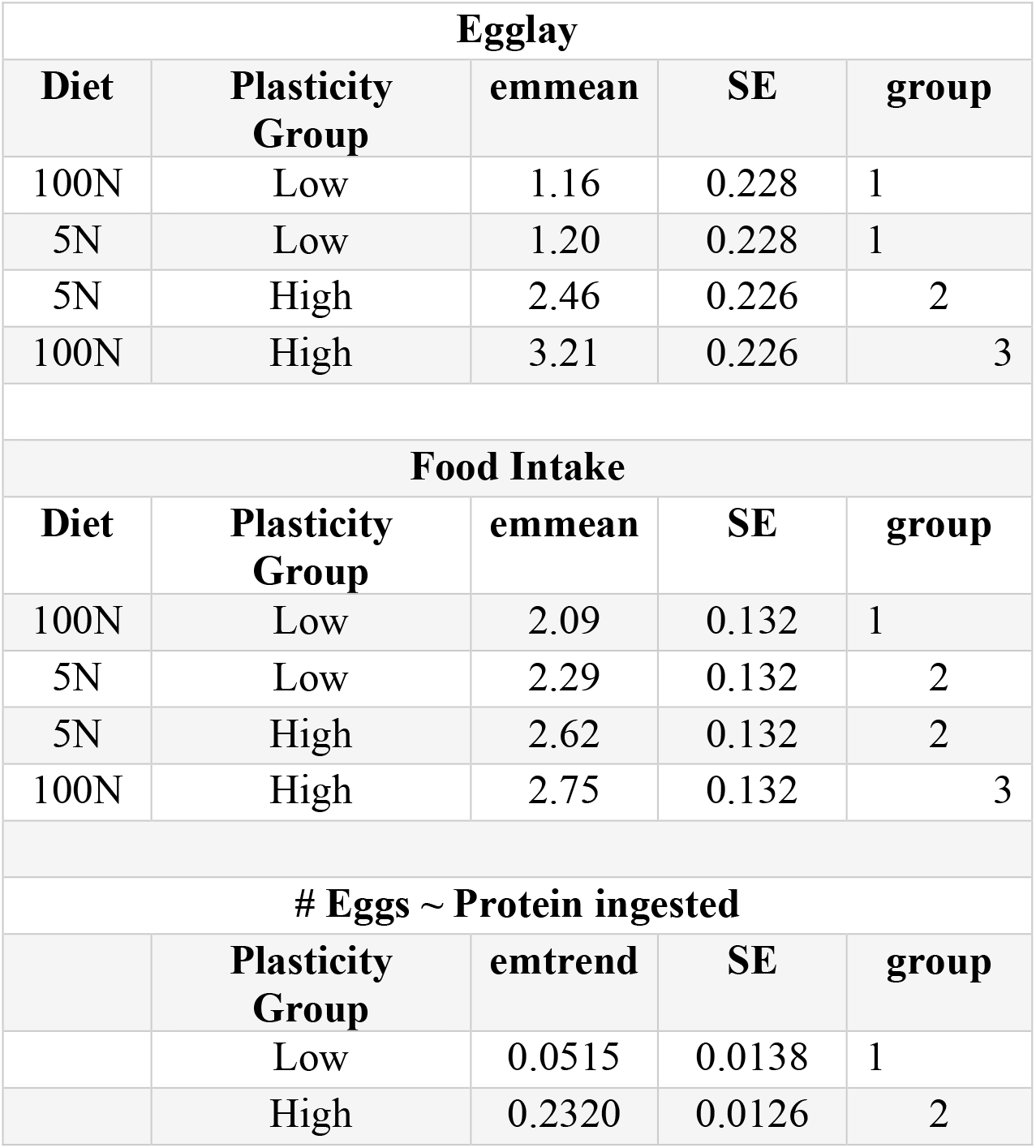
Means for number of eggs produced, and amount of food ingested and trends for amount of eggs produced by function of protein ingested. The group column represents statistically significant groups that differ from each other with p-value <0.05.

**Supplementary Figure 1:**
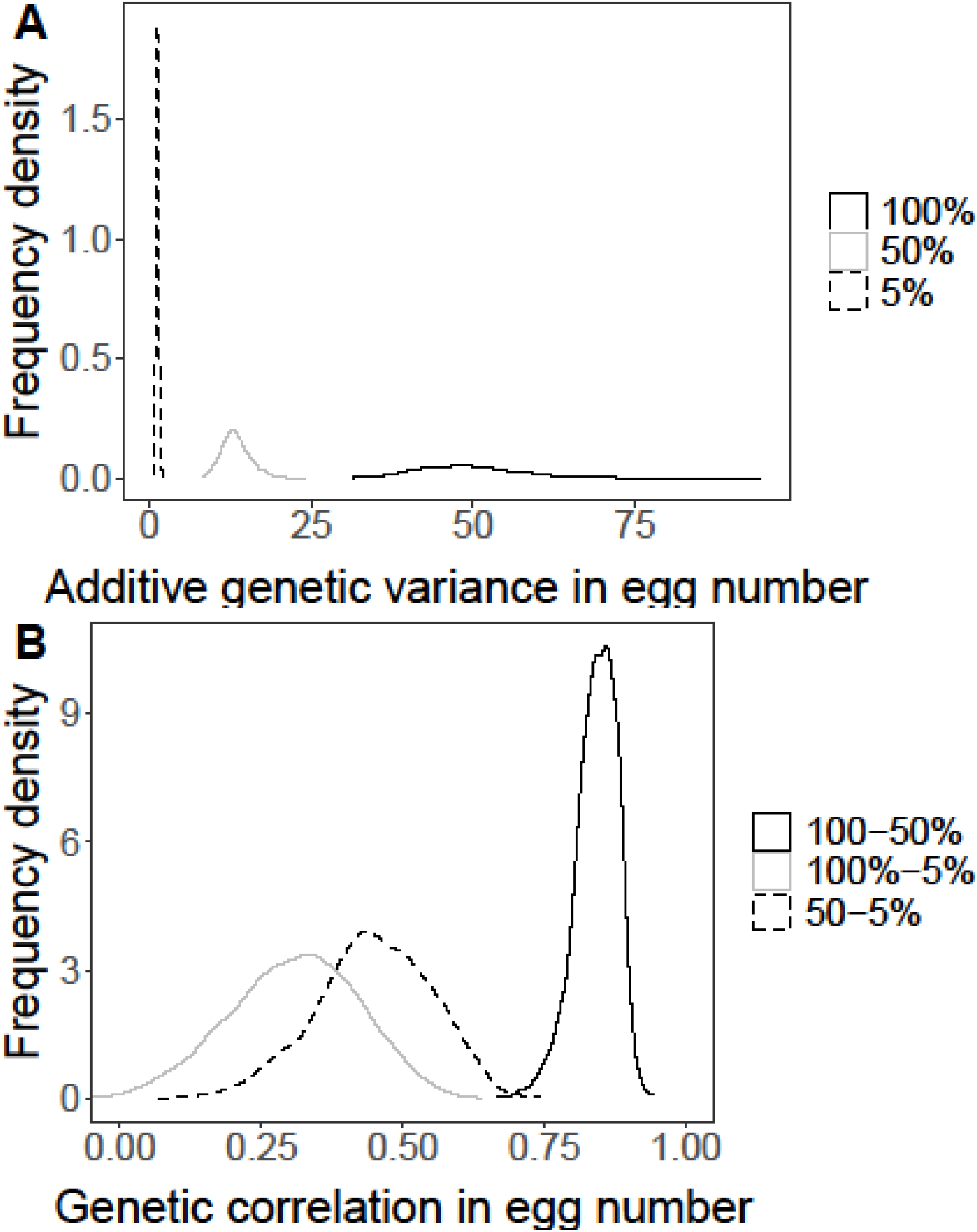
Density distributions of additive genetic variance and genetic correlations in egg number. A) Each line represents the density distribution of additive genetic variance in egg number for a particular diet. Solid black corresponds to the control diet, 100% yeast content, solid grey corresponds to 50% yeast content and dashed black line corresponds to the 5% yeast content diet. B) Each line represents the density distribution of genetic correlation in egg number for a particular pair of diets. Solid black corresponds to the comparison between control diet, 100% yeast content and 5% yeast content, solid grey corresponds to the comparison between 100% and 50% yeast content and dashed black line corresponds to the comparison between 50% and 5% yeast content diet.

