## Supplementary Tables and Figures for "Identifying the proximate mechanisms that generate variation in nutritional plasticity for fecundity in *Drosophila melanogaster*"

### Supplementary Figures and Tables

Supplementary Table 1 – Ovariole number does not differ among plasticity groups (high or low plasticity). Ovariole number was fit with a linear mixed effects model.

| Ovariole number | | | |
|---|---|---|---|
| **Term** | **Chi-square** | **Df** | **P-value** |
| **Plasticity** | 1.3248 | 1 | 0.2497 |
| **Model** | | | |
| **Number of ovarioles ~ Plasticity Group + (1\|Microtube) + (1\| Line)** | | | |

Supplementary Table 2: Means for number of eggs produced, and amount of food ingested and trends for amount of eggs produced by function of protein ingested. The group column represents statistically significant groups that differ from each other with p-value <0.05.

| Egglay | | | | |
|---|---|---|---|---|
| **Diet** | **Plasticity Group** | **emmean** | **SE** | **group** |
| 100N | Low | 1.16 | 0.228 | 1 |
| 5N | Low | 1.20 | 0.228 | 1 |
| 5N | High | 2.46 | 0.226 | 2 |
| 100N | High | 3.21 | 0.226 | 3 |
| **Food Intake** | | | | |
| **Diet** | **Plasticity Group** | **emmean** | **SE** | **group** |
| 100N | Low | 2.09 | 0.132 | 1 |
| 5N | Low | 2.29 | 0.132 | 2 |
| 5N | High | 2.62 | 0.132 | 2 |
| 100N | High | 2.75 | 0.132 | 3 |
| **# Eggs ~ Protein ingested** | | | | |
| | **Plasticity Group** | **emtrend** | **SE** | **group** |
| | Low | 0.0515 | 0.0138 | 1 |
| | High | 0.2320 | 0.0126 | 2 |

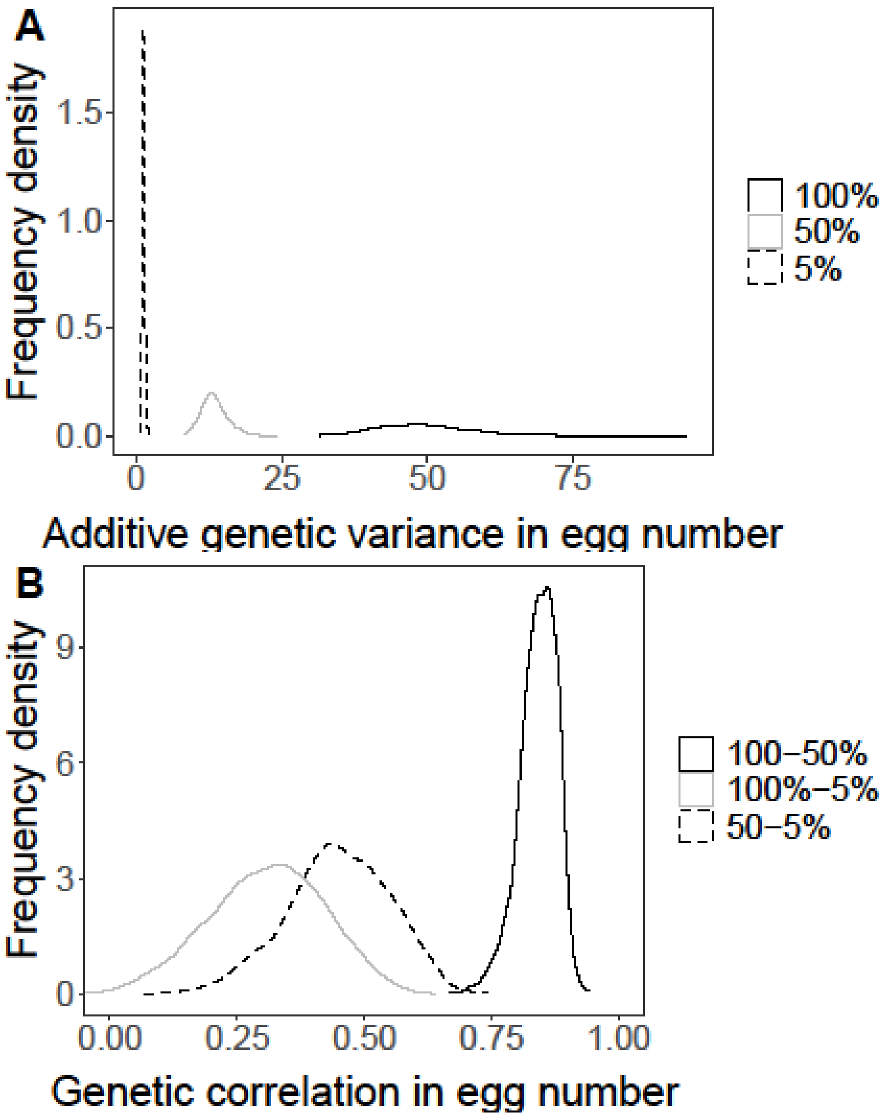

Supplementary Figure 1 – Density distributions of additive genetic variance and genetic correlations in egg number. A) Each line represents the density distribution of additive genetic variance in egg number for a particular diet. Solid black corresponds to the control diet, 100% yeast content, solid grey corresponds to 50% yeast content and dashed black line corresponds to the 5% yeast content diet. B) Each line represents the density distribution of genetic correlation in egg number for a particular pair of diets. Solid black corresponds to the comparison between control diet, 100% yeast content and 5% yeast content, solid grey corresponds to the comparison between 100% and 50% yeast content and dashed black line corresponds to the comparison between 50% and 5% yeast content diet.
